# Rapid development of an infant-derived HIV-1 broadly neutralizing antibody lineage

**DOI:** 10.1101/416032

**Authors:** Cassandra A Simonich, Laura Doepker, Duncan Ralph, James A Williams, Amrit Dhar, Zak Yaffe, Lauren Gentles, Christopher T Small, Brian Oliver, Vladimir Vigdorovich, Vidya Mangala Prasad, Ruth Nduati, D Noah Sather, Kelly K Lee, A Matsen Frederick, Julie Overbaugh

**Affiliations:** Division of Human Biology, Fred Hutchinson Cancer Research Center, Seattle, WA 98109, USA; Medical Scientist Training Program, University of Washington School of Medicine, Seattle, WA 98195, USA; Public Health Sciences Division, Fred Hutchinson Cancer Research Center, Seattle, WA 98109, USA; Department of Medicinal Chemistry, University of Washington, Seattle, WA 98195, USA; Department of Statistics, University of Washington, Seattle, WA 98195, USA; Department of Microbiology, University of Washington, Seattle, WA 98195, USA; Division of Basic Sciences, Fred Hutchinson Cancer Research Center, Seattle, WA 98109, USA; Center for Infectious Disease Research, Seattle, WA 98109, USA; Department of Pediatrics and Child Health, University of Nairobi, Nairobi, Kenya

## Abstract

HIV-infected infants develop broadly neutralizing plasma responses with more rapid kinetics than adults, suggesting the ontogeny of infant responses could better inform a path to achievable vaccine targets. We developed computational methods to reconstruct the developmental lineage of BF520.1, the first example of a HIV-specific broadly neutralizing antibody (bnAb) from an infant. The BF520.1 inferred naïve precursor binds HIV Env and a bnAb evolved within six months of infection and required only 3% mutation. Mutagenesis and structural analyses revealed that for this infant bnAb, substitutions in the kappa chain were critical for activity, particularly in CDRL1. Overall, the developmental pathway of this infant antibody includes features distinct from adult antibodies, including several that may be amenable to better vaccine responses.

---

Considerable efforts are spent on defining evolutionary pathways of broadly neutralizing antibodies (bnAbs) in HIV-1 infection under the premise that these pathways will help guide effective immunization strategies^1^. Particular emphasis has been placed on bnAb epitopes that are common in different individuals^2^, such as the V3-glycan region of HIV-1 envelope (Env)^3^, and much progress has been made toward characterizing the development of V3-glycan bnAbs in adults^4-7^. While the evolutionary pathway of adult bnAbs have been dissected to inform vaccine approaches^2^, significant challenges remain to be addressed for inducing bnAb responses by vaccination. One such challenge is that most adult bnAbs take years to develop as a result of a complex interplay between viral escape and antibody maturation^2^ that often leads to extensive somatic hypermutation (SHM), ranging from ∼6-29% (averaging ∼18%) for adult-derived V3-glycan bnAbs^4,6,8-13^. Additionally, the inferred germline precursors of a number of bnAbs lack detectable binding to recombinant HIV envelope and thus require the design of germline-targeting immunogens for vaccines^4,5,7,8,14,15^. Furthermore, some bnAbs are limited by autoreactivity^6,8,16^.

Recent studies reveal that infants and children develop bnAb responses at least as commonly, if not more frequently, than adults^17,18^ and that they do so rapidly, within 1-2 years post-infection^17^. To begin to characterize these early infant cross-clade neutralizing antibody responses, we previously isolated antibodies from an infant with a rapid and broad plasma nAb response. One infant-derived bnAb, BF520.1, demonstrated cross-clade neutralization breadth and targets the V3-glycan region of HIV Env. In contrast to most adult-derived bnAbs, BF520.1 has limited SHM (VH=6.6%, VK=5% nt)^19^. The infant immune system has many unique features compared to adults^20,21^ but the differences between infant and adult antibody development is not known. Here we identify the naïve antibody precursor of BF520.1, describe the subsequent evolution of this antibody lineage, and explore the structural basis for HIV binding for the first and currently only infant-derived HIV-specific bnAb that has been described.

## Results

### Ontogeny of the infant-derived bnAb BF520.1

Infant BF520 was infected with a clade A HIV virus, detectable at 3.8 months of age, and a broadly-neutralizing V3 glycan-directed monoclonal antibody (mAb) BF520.1 was isolated one year post-infection (pi)^19^ at 15 months of age. To infer the ontogeny of this bnAb, we performed next generation sequencing of the infant’s B cell repertoire on an available post-infection sample from 6 months pi (9 months of age). We inferred the naïve heavy chain (VH) and light chain (VK) ancestors of the BF520.1 mAb and identified all clonally-related sequences that descended from these ancestors (i.e. the clonal family of the bnAb) using the partis software package^22-24^, which is more accurate than previous methods but has not been previously applied to deep sequencing bnAb inference.

To illustrate where BF520.1 falls within the context of its clonal family, maximum likelihood (ML) phylogenetic trees were inferred for VH and VK clonal families (Fig. 1a,c). To reconstruct the most probable developmental routes taken by BF520.1 VH and VK sequences during the first six months of infection, we performed Bayesian phylogenetic and ancestral sequence inference analyses for each antibody chain. Importantly, this approach allowed us to rigorously estimate developmental route uncertainty because it accounts for phylogenetic uncertainty by considering many different phylogenies, which ML does not. We computationally validated that Bayesian lineage reconstruction correctly identified ancestral sequences for simulated antibody lineages with similar characteristics to that of BF520.1 97% of the time for ancestral sequences with posterior probability > 0.8, while dnaml and dnapars were less accurate (87% and 89%, respectively) (Supplementary Fig. 1). Therefore, inspired by Gong et al.’s approach for reconstructing influenza evolution^25^, we developed a software pipeline that uses Bayesian phylogenetics^26,27^ to reconstruct BF520.1 VH and VK lineages while retaining relative confidences for internal node sequences (Fig. 1b,d).

Using two replicates of NGS data, we selected the most probable routes of development for the BF520.1 heavy and light chains based on Bayesian lineage inference between infection and 6 months pi. We synthesized chosen lineage intermediates for further study: VH intermediates 1-3 (Int1-3_VH_) and VK intermediates 1-4 (Int1-4_VK_). Selection of these lineages relied on considering the most confident transitions (Fig 1b,d; darkest blue arrows) from the naïve sequence to the mature BF520.1 sequence in both NGS replicates. For the heavy chain, both replicates indicated that the G57D substitution occurred first and thus Int1_VH_ included this change. The Y32N amino acid substitution occurred next in both replicates and was therefore incorporated in Int2_VH_. The M34I and F114L substitutions were inferred in one of the two replicates, but the order was unclear, so both were added to Int3_VH_. For the kappa chain, both replicates supported L78M as the first mutation, so this substitution was added to create Int1_VK_. However, there was less agreement between replicates on the chronological order of the substitutions that followed. Both agreed that two substitutions (S30A and S67F) occurred prior to two later substitutions (S28N and T53S), so these steps were incorporated as Int2_VK_ and Int3_VK_. Finally, the A25T substitution was incorporated into Int4_VK_ because one of the two replicates indicated that this was a possible late substitution. Using this approach, we reconstructed highly probable routes of heavy and light chain antibody development within the first 6 months of HIV infection.

**Fig. 1:**
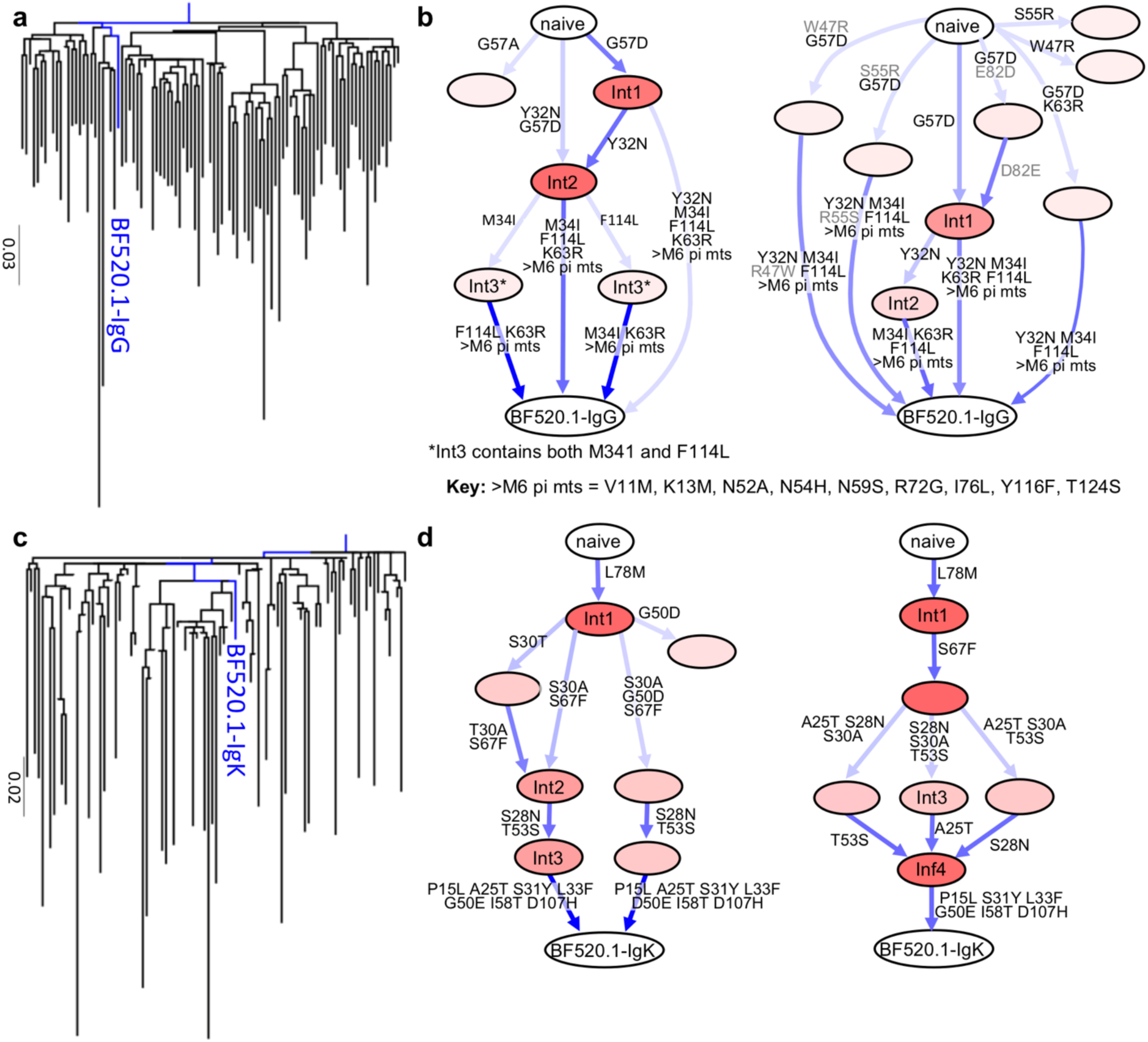
Ontogeny of the infant-derived bnAb BF520.1. a, c, Maximum likelihood phylogenetic relationships of (**a**) heavy and (**c**) light chain antibody gene variable regions. Trees display the inferred naïve ancestor (root), BF520.1 from M12 pi (blue), and representative clonal family member next generation sequencing (NGS) reads from M6 pi (black). Units for branch length estimates are nucleotide substitutions per site. **b, d**, Most probable routes of BF520.1 development: Bayesian BF520.1 clonal family phylogenies were sampled from an associated posterior distribution and then summarized to display relative confidences for internal node sequences. Resulting graphics display multiple possible lineages of amino acid transitions and their relative confidences for (**b**) heavy and (**d**) light chain development determined from two NGS technical replicates (left, right). Amino acid substitutions (arrows) connect the inferred naïve sequence to the mature BF520.1 sequence via reconstructed ancestral intermediate sequences (nodes). Labeled nodes were chosen as the most probable Bayesian lineage intermediate sequences. The red shading of nodes is proportional to the posterior probability that this ancestral sequence was present in the lineage. For a given node, the blue shading across arrows arising from that node is proportional to the corresponding transition probability. Transient mutations are labeled in grey.

### Increasing heterologous neutralization by the maturing BF520.1 heavy chain lineage

Antibodies were tested for HIV neutralizing activity against a panel of heterologous viruses that were selected from both the virus panel used to describe infant plasma neutralization breadth^17^ and the standardized “global panel”^28^ based on their ability to be neutralized by the mature BF520.1 antibody with an IC_50_<20 μg ml^-1^ ^19^. The inferred naïve mAb derived by pairing the inferred naïve VH and VK did not demonstrate neutralizing activity against any virus (Fig. 2a). To identify the VH substitutions gained within the first 6 months pi that were important for neutralization breadth, the naïve VH and Bayesian lineage intermediates (Int1-3_VH_) were paired with the mature VK and tested for HIV neutralizing activity. In contrast to the naïve mAb, the naïve VH paired with the mature VK (naïve_VH_mature_VK_) demonstrated cross-clade tier 2 neutralizing activity (Fig. 2a). The Int1_VH_mature_VK_ mAb had comparable activity to the naïve_VH_mature_VK_ mAb suggesting that the CDRH2 G57D substitution in this Int1_VH_ did not confer increased neutralizing activity (Fig. 2a,b). Subsequent VH intermediates demonstrated increasing heterologous, cross-clade neutralizing activity with affinity maturation. Notably, a Y32N substitution in the CDRH1 conferred increased neutralization breadth for Int2_VH_ and further cross-clade breadth and potency was observed in Int3_VH_ with FR2 M34I and CDRH3 F114L substitutions (Fig. 2a,b).

Because the BF520.1 clonal family sequences were collected from a midpoint sample (M6 pi), these sequences only informed the first 6 months of BF520.1 antibody development. Thus, for mutations that occurred in between M6 and M12 pi, we rationally incorporated amino acid substitutions in order of precedence: CDR mutations, CDR-adjacent framework (FR) mutations, then non-conservative FR mutations (Int4-6_VH_) (Fig. 2b). With the addition of N52A in the CDRH2 of Int4_VH_, the antibody gained breadth and potency comparable to the mature mAb BF520.1 (Fig. 2a,b).

Previous studies inferred intermediate sequences of antibody lineages from single ML and parsimony (Pars) phylogenies^6,29-31^ and thus did not consider relative confidence as we do with our Bayesian approach. Given this precedent, we compared our results using Bayesian methods against corresponding single ML (Supplementary Fig. 2a) and Pars phylogenies (Supplementary Fig. 2c) to infer BF520.1 intermediates. Interestingly, the VH intermediates inferred from the ML phylogeny included substitutions that were not found in either the naïve or mature mAbs, indicating that they were likely erroneous. The Bayesian lineages also included substitution reversions, but these paths consistently had lower posterior probabilities than paths lacking these artifacts (Fig. 1b,d). When present, these substitutions decreased neutralizing activity, which is apparent when comparing ML Int3_VH_ and ML Int4_VH_ (Supplementary Fig. 2a,b). These findings suggest that the Bayesian antibody lineage determination that incorporated relative confidence over a number of possible lineage pathways prevented the inference of artifactual transient mutations in antibody intermediates. Regardless, our studies of the ML and Pars lineage intermediates led to similar conclusions about the importance of heavy chain substitutions: CDRH1 Y32N and CDRH2 N52A substitutions resulted in increased neutralization breadth (Supplementary Fig. 2a-d). The contribution of the Y32N CDRH1 substitution was particularly apparent when comparing Pars Int3_VH_, which lacks Y32N and Pars Int4_VH_ (contains Y32N) (Supplementary Fig. 2c,d).

Overall, these data demonstrate increasing heterologous neutralizing activity with affinity maturation and show that the heavy chain CDRH1 Y32N and CDHR2 N52A substitutions are important for the neutralization breadth of BF520.1.

**Fig. 2:**
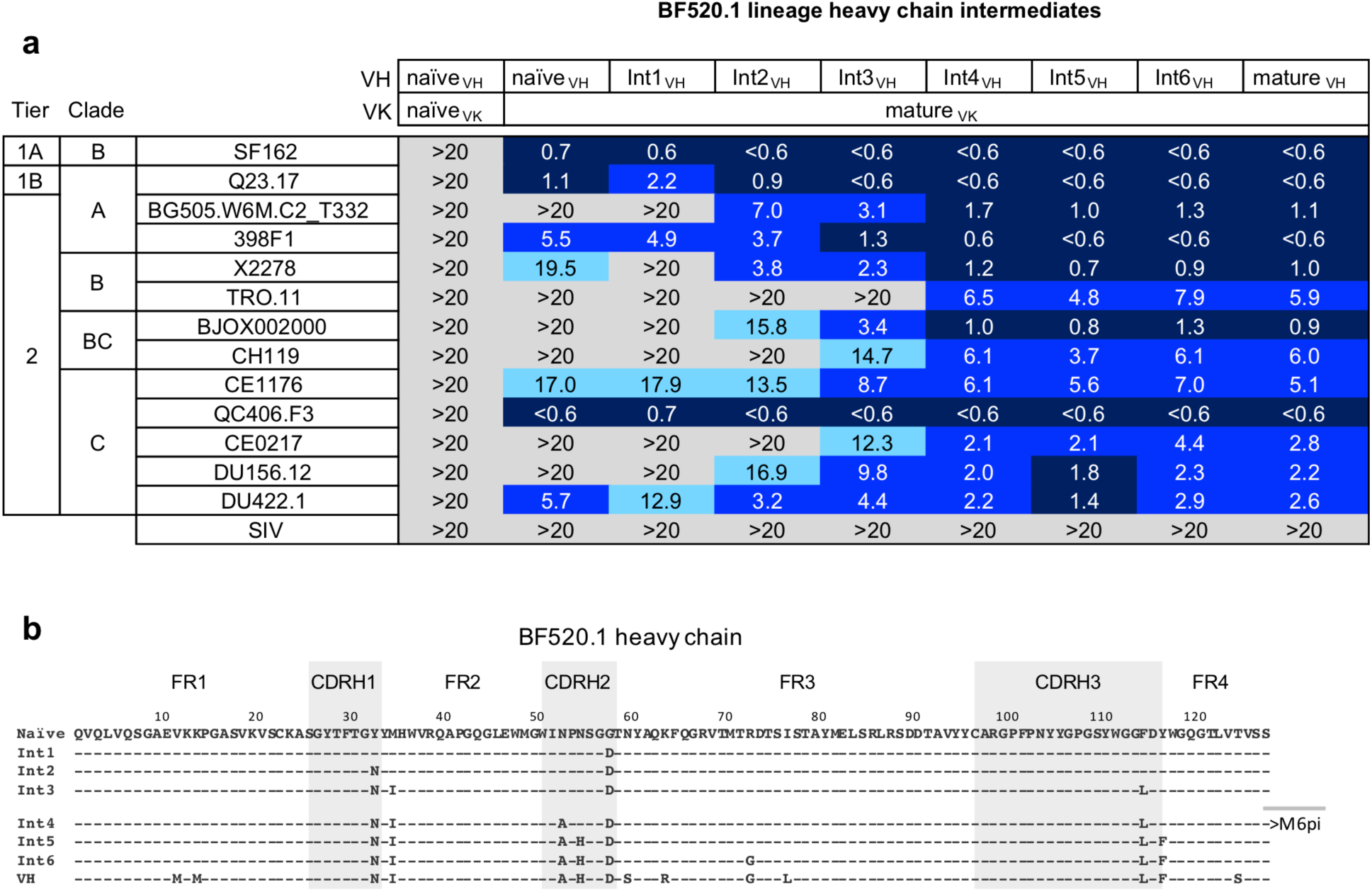
Development of heterologous neutralization by the maturing BF520.1 heavy chain lineage. a, Neutralization of panel viruses by BF520.1 inferred lineage intermediates. The top rows of the table (VH and VK) show the origin of the antibody chain sequence used indicated by Int#_VH_ with # indicating the progression of intermediates in the lineage. These were paired with the indicated kappa chain, in most cases the mature kappa chain (mature_VK_). The panel viruses are shown to the left, with the tier, clade and name indicated. IC_50_ values (μg ml^-1^) represent an average of two to three independent experiments performed in duplicate. IC_50_ values are color-coded with darker shades indicating more potent neutralization. Grey indicates that 50% neutralization was not achieved at the highest mAb concentration tested. **b**, Amino acid alignment of BF520.1 naïve, Bayesian (0-6 months pi) and rationally inferred (6-12 months pi) lineage intermediates and mature heavy chain sequences. Intermediates designated as >M6 pi were rationally inferred.

### Contribution of kappa light chain maturation to HIV binding and neutralization

Given the surprising finding that the naïve VH paired with the mature VK (naïve_VH_mature_VK_) demonstrated cross-clade tier 2 HIV neutralizing activity and the naïve mAb did not (Fig. 2a), we also examined the evolution of VK in relation to binding of HIV Env and neutralization breadth. The cross-paired mature and naïve heavy and light chains (naïve_VH_mature_VK_ and mature_VH_naïve_VK_) both bound the BG505 SOSIP trimer, which is an Env that is sensitive to BF520.1 neutralization. The naïve_VH_mature_VK_ mAb demonstrated stronger binding to the BG505.SOSIP.664 trimer (K_D_=<0.001nM) (Fig. 3a) than the converse mature_VH_naïve_VK_ mAb (K_D_=8.95nM) (Fig. 3b). In contrast to the heterologous neutralizing activity seen for the naïve_VH_mature_VK_ mAb, the mature_VH_naïve_VK_ mAb neutralized only tier 1A SF162 variant but not tier 2 viruses (Fig. 3b). These data suggest that maturation in VK is necessary for BF520.1 heterologous neutralization breadth.

To identify which mutations in VK contributed to increased neutralizing activity, VK Bayesian lineage intermediates (Int1-4_VK_) and subsequent rationally-inferred intermediates (Int5-7_VK_) were paired with the mature VH and tested for cross-clade tier 2 neutralizing activity. Overall, VK lineage intermediates demonstrated increasing breadth with maturation (Fig. 3c,d). A dramatic jump in neutralization breadth was demonstrated by Int2_VK_, which contains CDRL1 S30A and FR3 S67F substitutions (Fig. 3c,d). Although S67F was observed prior to the S30A substitution in the ML and Pars VK lineages, it did not increase neutralization breadth (Supplementary Fig. 3a-d), suggesting that the S30A CDRL1 substitution alone enables the observed heterologous neutralization breadth. Further augmentation of potency and breadth was conferred by substitutions observed in Int3_VK_ (CDRL1 S28N and CDRL2 T53S). The T53S substitution was observed prior to S28N in the ML VK lineage and it did not increase breadth but did confer a slight increase in potency. Int4_VK_ (FR1 A25T) had a very modest increase in activity. Importantly, Int5_VK_ (CDRL1 S31Y and CDRL2 G50E) neutralized all viruses that were neutralized by the mature antibody and Int6_VK_ (FR2 L33F) reached potency comparable to BF520.1 (Fig. 3c,d).

While there are multiple mutations in and around CDRL1, the BF520.1 VK does not contain CDRL3 mutations, based on the inferred naïve sequence. These data suggest that kappa chain mutations in and around the CDRL1 are important for BF520.1 heterologous neutralization breadth.

**Fig. 3:**
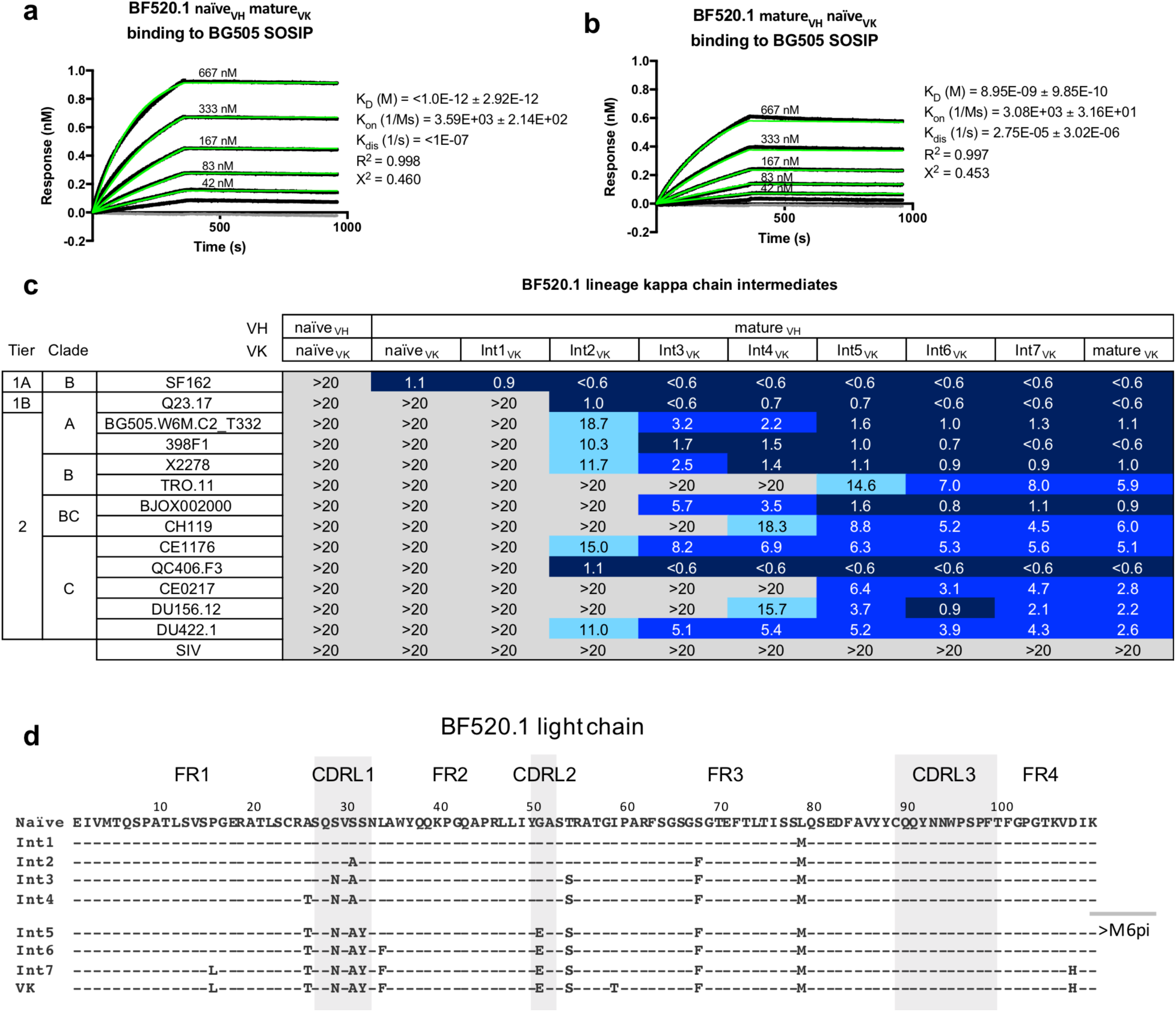
Contribution of kappa light chain maturation to HIV binding and neutralization. a,b, BLI representative reference-subtracted sensorgrams for each interaction between the BG505.SOSIP.664 (ligand) and BF520.1 naïve_VH_mature_VK_ (**a**) and mature_VH_naïve_VK_ (**b**) (analyte). IgG concentrations ranged from 667 to 42 nM. The gray lines show 0 μM IgG. K_D_, K_on_, and K_dis_ are shown from best fitting (green lines) to a 1:2 bivalent analyte model of binding. **c**, Neutralization of panel viruses by BF520.1 inferred lineage intermediates. Layout is as described for Fig. 2. **d**, Amino acid alignment of BF520.1 naïve, Bayesian (0-6 months pi) and rationally inferred (6-12 months pi) lineage intermediates and mature kappa chain sequences. Intermediates designated as >M6 pi were rationally inferred.

### Cross-clade neutralization with limited SHM in paired intermediates

To define the minimal combination of changes in VH and VK that are needed for breadth, intermediates were paired together using percentage SHM to assign probable pairings. Tier 1A neutralizing activity was observed following just two amino acid substitutions in VH (Int2_VH_) and three in VK (Int2_Vk_) (Fig. 4) with the additions of the VH Y32N and VK S30A being important for the neutralizing activity. Cross-clade heterologous neutralization was achieved with the addition of the VH M34I and VK S28N in the Int3_VH_Int3_Vk_ mAb, which had a total of four VH and five VK amino acid substitutions or 1.3% and 1.8% SHM at the nucleotide level in VH and VK, respectively. Int4_VH_ paired with Int4_VK_ neutralized the majority of the viruses (9/13; 69%) that are neutralized by mature BF520.1 with only 1.8 VH and 2.1% VK SHM. Further increased neutralization was conferred by single additional substitutions in both VH and VK: N52A (Int4_VH_) and A25T (Int4_VK_). The breadth of BF520.1 was reached with 1.8% VH and 3.4% VK SHM (2.5% overall SHM) for Int4_VH_Int5_VK_. Finally, the Int5_VH_Int6_VK_ mAb reached the breadth and potency of mature BF520.1 with only 2.4% VH and 3.7% VK SHM (3.0% overall SHM) (Fig. 4). Similar observations were made using the ML (Supplementary Fig. 4a) and Pars (Supplementary Fig. 4b) lineages.

**Fig. 4:**
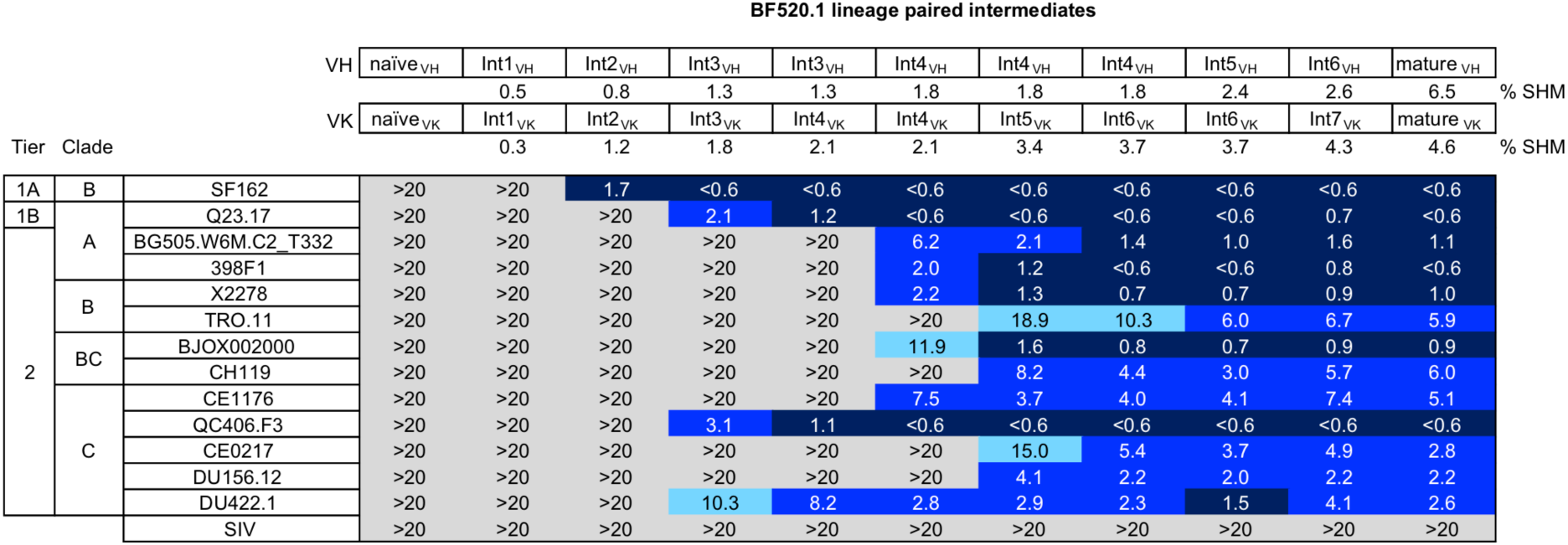
Neutralization properties of paired intermediates of VH and VK. The top rows show the intermediate in the lineage tested for both VH and VK chains as shown in Fig. 2b and 3d. Below each the % SHM is indicated. The virus panel and figure layout is as described in the legend to Fig. 2a.

### Cryo-EM structure of the BG505.SOSIP.664 trimer in complex with BF520.1 Fab

Single particle cryo-electron microscopy was used to gain insight into the interaction of BF520.1 Fab and BG505.SOSIP.664 trimer, which is a clade A transmitted envelope variant from a Kenyan infant who was in the same cohort as BF520^32,33^. The resulting 4.8Å map (Fig. 5 and Supplementary Fig. 5) revealed features consistent with the resolution estimate including helices, beta-sheet structure as well as bulky densities for glycans that protrude from glycosylation sites, including N332 and N301, which are critical features for BF520.1 activity^34-37^ (Supplementary Fig. 5e).

The positioning of the CDRH loops in our structural model indicates that they make multiple contacts with both protein and N-linked glycans (Fig. 1b). The CDRH1 loop is in close proximity to the base of the gp120 V3-loop, with N32 oriented towards the conserved GDIR sequence, consistent with increased neutralization observed when asparagine is introduced at position 32 on the CDRH1 loop (Y32N substitution). Adjacent residues of CDRH1 as well as the CDRH2 loop are located proximal to the N301 glycan. It is unclear if residue 52 (N52A substitution) in CDRH2 mediates direct contact between either the GDIR sequence or N301 glycan; however, it is plausible that substitution for alanine could influence loop positioning such that residues H54 and S55 would interact with the N301 glycan. The relative orientation of the CDRH3 loop suggests it may interact with the V3 GDIR sequence as well as the N332 glycan, though given variability in CDRH3 positioning between homology models, the contacts are less certain for this loop.

The structure of the Env trimer in complex with BF520.1 supports a role for the BF520.1 VK in neutralization through extensive contacts between the CDRL1 loop and the N332 glycan. Based upon the BF520.1 homology model, N28, A30, and Y31, which were implicated in nAb breadth, are positioned directly adjacent to the N332 glycan (Fig. 5c), although we cannot unambiguously identify the position of specific residues. L33F and G50E mutations contributed to increased potency, but from the structure it is unclear whether these residues could directly contact the N332 glycan or protein surface^38,39^; they may instead alter local paratope structure in a more indirect fashion. The position of CDRL3 loop suggests it likely does not interact with the BG505 trimer or key glycans, which may explain why it does not contain mutations that contribute to BF520.1 breadth.

**Fig. 5:**
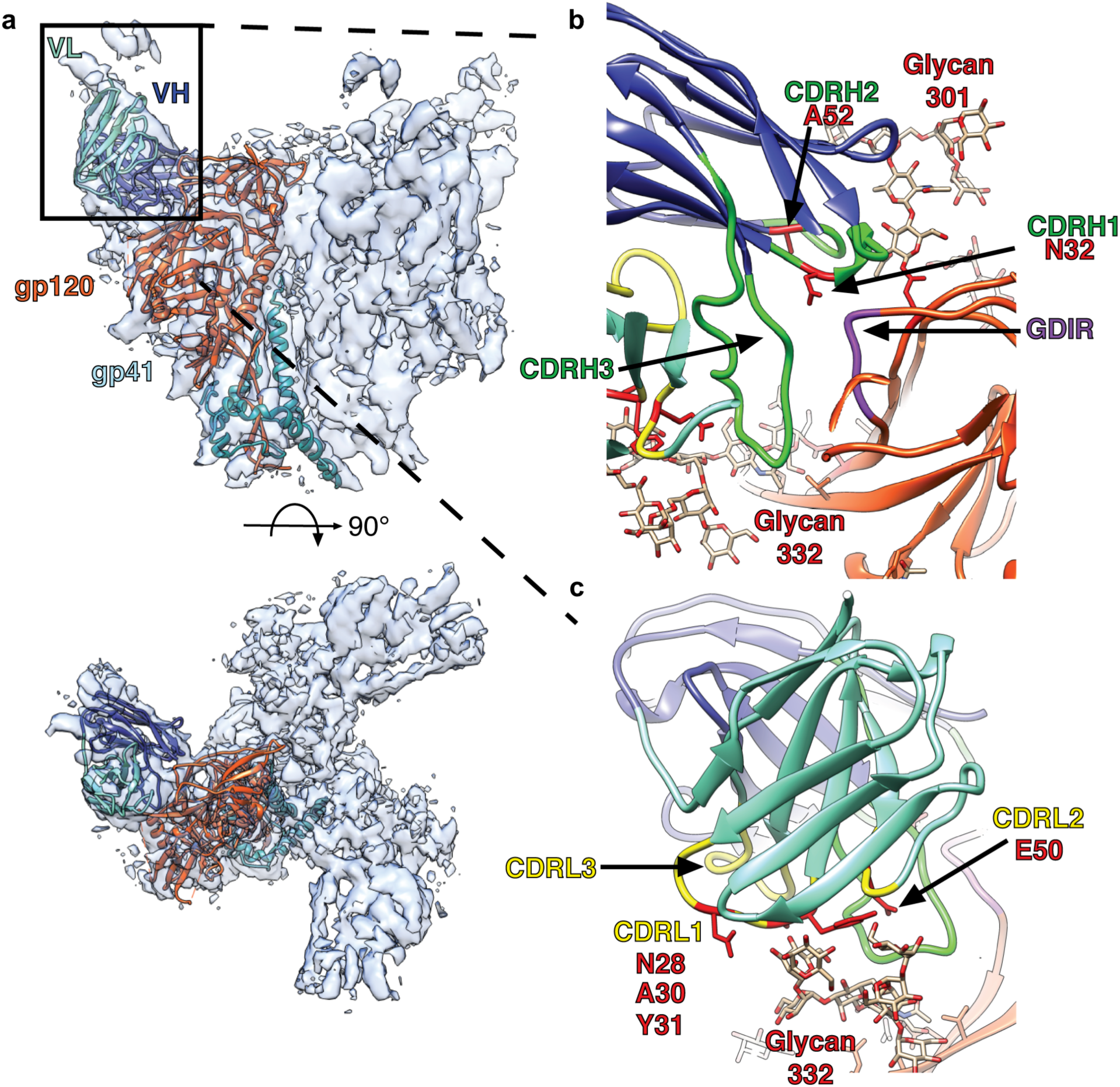
Cryo-EM reconstruction and model of the BG505.SOSIP.664 trimer in complex with BF520.1 Fab. a, Side (above) and top (below) views of the cryo-EM reconstruction and structural model. A single monomer consisting of gp120 (orange) and gp41 (dark cyan), and the BF520.1-Fv variable heavy chain (dark blue) and variable light chain (aquamarine) are highlighted. Glycans removed for clarity. The BG505.SOSIP trimer structure 5ACO.pdb docked into the new EM density map with a correlation score of 0.8602^34^. A global search yielded a preferred placement of the BF520.1 Fv model into the Fab density with a correlation score of 0.8513 (Supplementary Figure 5g). **b**, Expanded view of the VH domain and gp120. Shown are the conserved gp120 GDIR sequence (purple), glycans N332 and N301, and CDRH loops (green). Mutations that confer potent neutralization (red) are indicated. **c**, Expanded view of the VL domain and the N332 glycan. CDRL loops (yellow) and mutations that conferred potent neutralization (red) are highlighted.

### BF520.1 naïve and lineage members bind HIV Env trimer

Though the inferred naive ancestor of BF520.1 did not demonstrate neutralizing activity, it weakly bound to BG505.SOSIP.664 trimer (Fig. 6a) and this binding was dependent on increasing mAb concentration (Fig. 6b). Qualitatively, we observed increased binding with affinity maturation (Fig. 6a) with the most significant increase in binding demonstrated by Int2_VH_ Int2_VK_ and more subtle increases by later intermediates. Mutations acquired by Int4_VH_4_VK_ enabled similar binding to Env as the mature BF520.1. Despite clear binding to the BG505 SOSIP trimer by Int2_VH_2_VK_, Int3_VH_3_VK_, and Int3_VH_4_VK_, these intermediates were not able to neutralize the BG505 pseudovirus (Fig. 4 and 6a). These data indicate that the BF520.1 naïve mAb had the potential to recognize HIV Env.

Our approach to inferring the naïve ancestor using replicate sequence data differed from previous studies, which only used single replicates of NGS data^6,29,30^. To compare the two methods, we also inferred the BF520.1 naïve ancestor using just the first replicate of antibody sequence data. This approach resulted in uncertainty in the naïve nucleotide sequences at three positions (Supplementary Fig. 6a,b) while there was not uncertainty using replicate datasets. When tested, none of the alternative inferred naïve mAbs demonstrated neutralization (Supplementary Fig. 6c), nor did they bind trimer (Supplementary Fig. 6d), which contrasts the ability of the more probable replicate-inferred naïve mAb. We also found differences in the autoreactivity of the inferred naïve mAbs: single dataset-inferred mAbs demonstrated weak autoreactivity (Supplementary Fig. 6e), whereas this was not observed for the more probable replicate-inferred naïve mAb. These data demonstrate that uncertainty in the inference of bnAb precursors can result in significant differences in HIV Env binding and autoreactive properties.

**Fig. 6:**
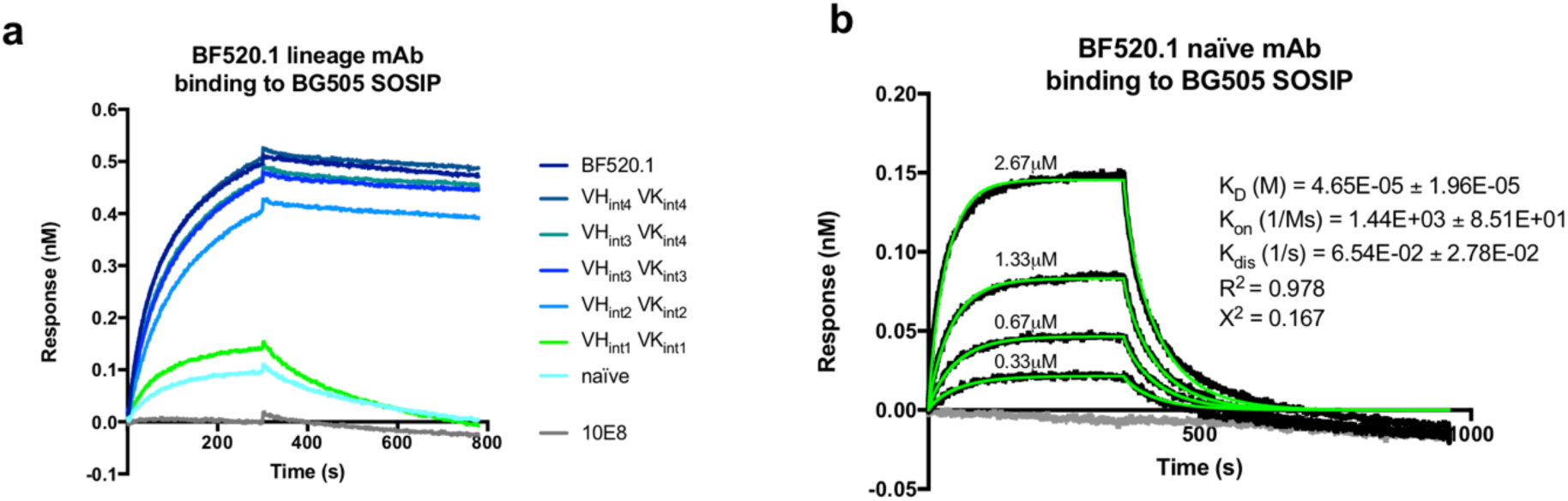
BF520.1 naïve and lineage binding to HIV Env trimer. a, BF520.1 paired lineage intermediates (ligand) binding to the BG505.SOSIP.664 (analyte) measured by BLI. 10E8 is a negative control as the SOSIP trimer does not contain the targeted MPER epitope. Data are representative of two independent experiments. **b**, BLI binding analysis of varying concentrations of BF520.1 naïve antibody (analyte) binding to BG505.SOSIP.664 (ligand). The gray line shows 0 μM IgG. K_D_, k_on_, and k_dis_ from best fitting (green lines) to a 1:2 bivalent analyte model of ligand:analyte binding are shown.

### Increasing autologous neutralization by the BF520.1 lineage

BF520.1 does not neutralize the transmitted autologous virus but it does neutralize variants isolated from 6 months of age (M6), which is ∼2.2 months after infection was detected^19^. Similar to the mature mAb BF520.1, the naïve and early lineage intermediates were not able to neutralize a transmitted autologous envelope variant (Supplementary Fig. 7). Likewise, the earliest paired intermediates failed to neutralize the M6 variants. Lineage member Int4_VH_4_VK_ neutralized BF520.M6P.I1 with lower potency compared to BF520.1 (5.8 vs. 1.1 μg ml^-^ 1, respectively). Additionally, Int4_VH_5_VK_ and Int4_VH_6_VK_ mAbs demonstrated stepwise increases in neutralization of the autologous viruses from ∼2.2 months pi, further highlighting the importance of the kappa chain contribution to autologous neutralizing activity. Substitutions in the CDRL1 and CDRL1-proximal regions were particularly impactful for autologous neutralization: CDRL1 S31Y, CDRL2 G50E and FR2 L33F (Supplementary Fig. 7).

## Discussion

The complex developmental pathways of bnAbs in HIV-infected adults have been carefully examined to guide immunogen design targeting key precursors in antibody evolutionary pathways^2^. In many cases, this approach is complicated by the observation that the inferred naïve progenitor of the bnAb does not bind HIV Env, presenting challenges to stimulating the lineage^15^. Taking advantage of the rapid development of HIV bnAb responses in infants, here we discovered an evolutionary pathway that starts with a naïve antibody that is capable of binding HIV Env trimer and requires as little as 3% SHM to achieve the full cross clade breadth of this lineage. Remarkably, this antibody lineage relies on changes in both the heavy and light chains, mostly outside of the CDRH3, to develop HIV specificity and breadth. Thus, this antibody developmental pathway allows for two key aspects of vaccine design that were stumbling blocks for many adult bnAbs pathways: the HIV antigen can engage the naïve B cell and a level of SHM that is easily within reach by vaccination^40^.

V3-glycan targeting bnAbs are of high interest because they are common and not restricted by certain germline genes^4,6,8,10,12,19^. BF520.1 is of particular interest because it developed more rapidly than adult V3-bnAbs and achieved broad and potent capability with as little as 3.0% SHM (2.4% VH and 3.7%VK SHM). Other known adult V3-glycan bnAbs required rare mutation events to occur^4,7^ and/or higher levels of SHM^6^ compared to BF520.1^41^. The highly impactful N52A substitution in the BF520.1 heavy chain required one nucleotide mutation in an AID hotspot site and one mutation in a neutral site, but the improbability of these mutations was not a barrier to rapid bnAb development in this infant. This indicates that vaccines may be able to stimulate similar mutations in reasonable timeframes.

Interestingly, mutations important for increasing BF520.1 functional activity were primarily found in the heavy chain CDRH2 and light chain CDRL1, with little-to-no contribution from the CDRH3 and CDRL3. The Cryo-EM structural model indicated that CDRH2 and CDRL1 residues appear to contribute to the BF520.1 paratope by mediating contacts with the conserved V3-glycan site, namely the archetypal features of V3-targeting bnAbs: GDIR sequence and glycans at position 301 and 332^42^. In contrast to the position of CDRL3, located too distant to make meaningful contacts with the trimer, the structure suggests that CDRH3 may interact with both the V3 GDIR sequence and N332 glycan; however, the lack of substitutions in the loop during SHM suggests a likely minor role for CDRH3 in BF520.1’s maturation. This contrasts many HIV bnAbs where major determinants of breadth reside in the CDR3 regions, particularly in VH^44^.

We identified that maturation in the BF520.1 VK was particularly important for HIV Env binding and neutralization, demonstrated by the increased heterologous breadth observed when the mature VK was paired with the naïve VH and the five amino acid substitutions in and around CDRL1 that contribute to neutralization breadth. The structural findings support this critical role for variable light chain development in the potent and broad neutralization capacity of BF520.1. With few exceptions^7^, functional studies have largely focused on the contribution of VH to heterologous neutralizing activity^4,6^ despite previous structural studies showing that adult-derived V3-glycan bnAb light chains contact the N332 glycan and conserved GDIR motif^10,43^. Thus, our studies highlight the importance of considering both the heavy and light chain in the development of HIV-specific bnAbs and the fact that the light chain can potentially be harnessed to develop vaccine approaches that elicit such bnAbs with relatively little SHM.

While we see neutralization of heterologous Env variants by the early BF520.1 lineage intermediates, autologous neutralization is observed only with later stage intermediates (Int4_VH_Int4_VK_ and beyond) against BF520 autologous Env variants isolated at ∼2 months pi. While CDRH2 and CDRL1 mutations are important for heterologous breadth, autologous neutralization was conferred by additional kappa chain mutations in and around the CDRL1. The observation that the earlier intermediates are HIV-specific but do not neutralize the autologous virus suggests that the virus at ∼2 months pi may have escaped neutralization by the early lineage. We do not know whether the naïve or early lineage mAbs were able to bind autologous viruses at the time of infection or at ∼2 months pi. The BF520.1 naïve mAb and early lineage intermediates do, however, bind the heterologous clade A BG505.SOSIP.664 trimer.

This binding of the naïve ancestor of BF520.1 to recombinant HIV Env contrasts with many adult bnAb precursors^15^, including some V3-glycan bnAbs^4,7^. The affinity of the BF520.1 naïve mAb for the Env trimer is within the range of affinities observed for VRC01 class naïve precursors binding to a monovalent germline-targeting antigen^45^. This study raises the possibility that an immunization strategy to elicit a BF520.1-like response could use the BG505.SOSIP.664 or an optimized target to initiate the response and then could be boosted with BF520 Env trimer from ∼2 months pi to drive affinity maturation.

To identify appropriate vaccine immunogens that might stimulate bnAb precursors, it is critical that inferred bnAb precursor heavy and light chain sequences are as accurate as possible. Importantly, accurate naïve inference enables more accurate phylogenetic inference of intermediate sequences. By using replicate NGS data, we resolved uncertainties in the inferred naïve sequences that impacted the overall characterization of our antibody lineage. We are confident in our BF520.1 naïve inference for three reasons: 1) we used the entire clonal family to infer the naïve sequence along with more accurate clonal family clustering, which together increase confidence especially within the CDR3^22,23^, 2) we used per-sample germline inference^24^, which mitigates gene assignment errors and expands our ability to accurately infer antibody ontogenies in non-European subjects whose antibody gene usage patterns are less well documented, and 3) BF520.1 had very low SHM compared to other bnAbs, which alleviated the difficulty of naïve inference due to a reduced expected number of incorrect bases. Notably, our most probable replicate-inferred naïve mAb measurably bound HIV and did not demonstrate autoreactivity. Conversely, the single-dataset inferred naïve mAbs that contained uncertainties displayed autoreactivity, which is misleading because this is indeed a feature of many adult bnAbs^16^ and some V3-glycan bnAbs^6,8^. These discrepancies highlight the potential imprecision of naïve antibody inference due to insufficient quality or depth of sequencing data and suggest that our newer approaches designed specifically for antibody lineage analysis should be used.

The finding that infants develop broad and potent HIV bnAb responses has raised the question of whether HIV vaccine efforts should focus on infants^20^. Indeed, pediatric immunizations for the prevention of viral infections is a practical strategy because infants and children have regular contact with the healthcare system. Understanding the ontogeny of HIV neutralizing antibody responses in the pediatric population is necessary to facilitate the rational design of such vaccine strategies. Given that the BF520.1 naïve ancestor binds HIV Env, that very little SHM is required for this V3-directed antibody lineage to demonstrate heterologous cross-clade breadth, and that these antibodies lack unusual or rare features, the BF520.1 lineage should be considered an attractive template for vaccine design. These studies suggest that the setting of mother-to-child transmission or the infant immune system may be uniquely poised to develop bnAb responses rapidly.

## Acknowledgements

We thank the participants and staff of the Nairobi Breastfeeding Trial. We thank Will DeWitt for help with sequence data analysis and Vrasha Chohan for assistance with neutralization assays. This work was supported by NIH grants R01 AI12096, R01 GM113246, U19 AI117891, and F30 AI122866. The research of Frederick Matsen was supported in part by a Faculty Scholar grant from the Howard Hughes Medical Institute and the Simons Foundation. Amrit Dhar was supported in part by an NSF IGERT DGE-1258485 fellowship. We thank the Arnold and Mabel Beckman Foundation and Washington Research Foundation for their support of the University of Washington Cryo-EM Center.

## Author Contributions

J.O. conceived the study. J.O., C.A.S., L.D., J.A.W., K.K.L., F.A.M., N.S., V.M.P., A.D., and D.R. all contributed to the design of the study. C.A.S., L.D., J.A.W., Z.Y., and L.G., performed experiments. R.N. developed the cohort and collected samples. All authors contributed to the analysis and interpretation of data. C.A.S., L.D., and J.O. wrote the manuscript with input from all authors.

---

**Supplementary Fig. 1:**
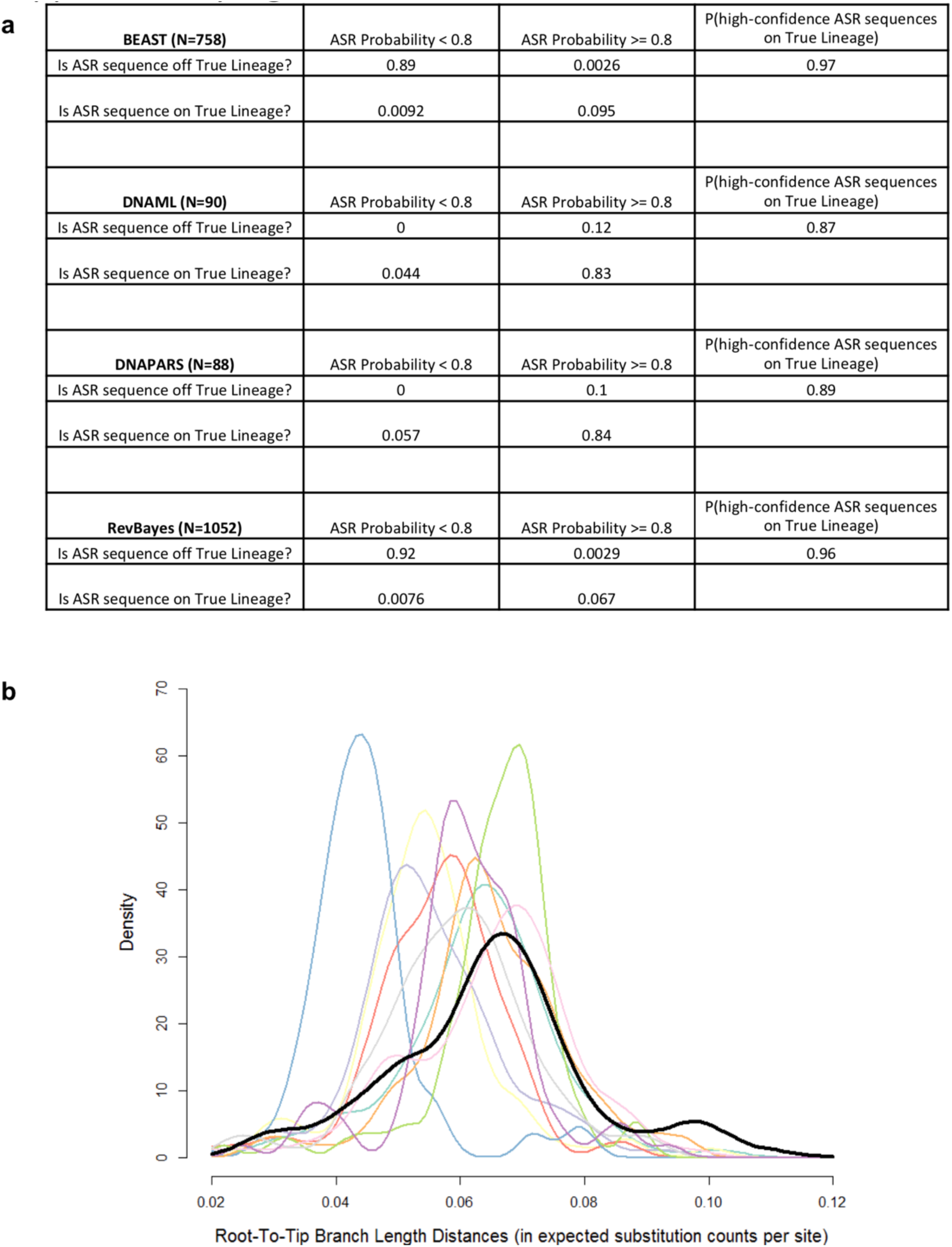
Accuracy of Bayesian lineage reconstruction for BF520.1-like simulated sequence alignments. a, Comparison of BEAST, dnaml, dnapars, and RevBayes methods for antibody lineage reconstruction based on simulated antibody phylogenies. True Lineages were defined as the naïve-to-seed lineages on simulated clonal family trees. P(X|Y) means the probability of X given Y. **b**, Comparison of the root-to-tip branch length distance distribution for the BF520.1 clonal family (in black) against the corresponding distributions for 10 independently simulated clonal families (in lighter colors), showing concordance between the simulations and the observed distribution. To compute the distribution for the BF520.1 family, we obtained 10,000 independent tree samples from RevBayes and computed the median root-to-tip branch length distance for each sequence in the clonal family. Each of the 10 simulated clonal families were obtained using the same simulation parameters.

**Supplementary Fig. 2:**
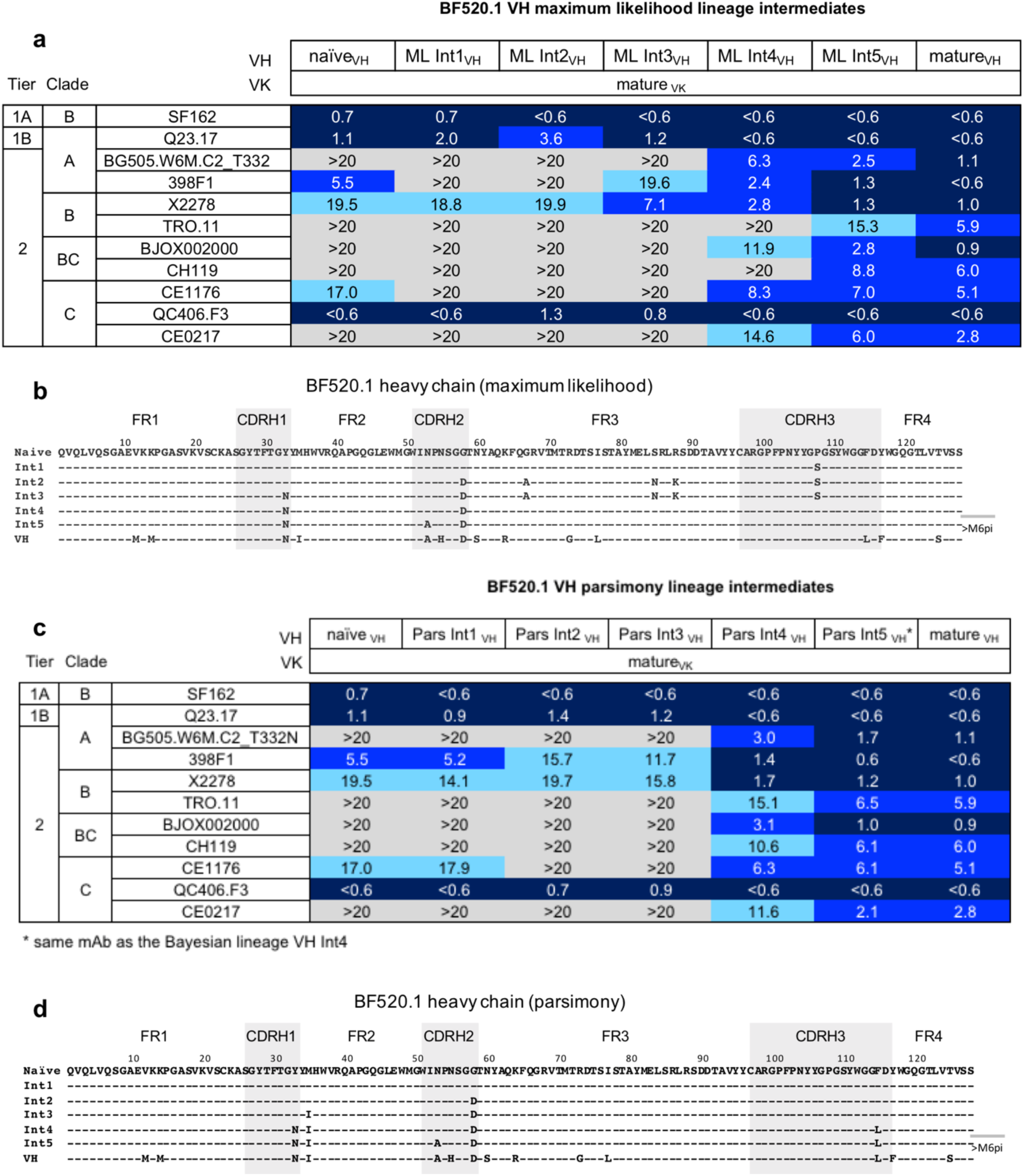
Increasing heterologous neutralization by the maturing BF520.1 heavy chain lineage. BF520.1. a,c, Maximum likelihood (**a**) and parsimony (**c**) lineage heavy chains paired with the mature kappa light chain, mAb neutralization of viruses. IC_50_ values (μg ml^-1^) are an average of two to three independent experiments performed in duplicate. IC_50_ values are color-coded with darker shades of blue indicating more potent neutralization. Grey indicates that 50% neutralization was not achieved at the highest mAb concentration tested. **b**,**d**, Amino acid alignments of maximum likelihood (**b**) and parsimony (**d**) heavy chain lineage intermediates. Intermediates designated as >M6 pi were rationally inferred.

**Supplementary Fig. 3:**
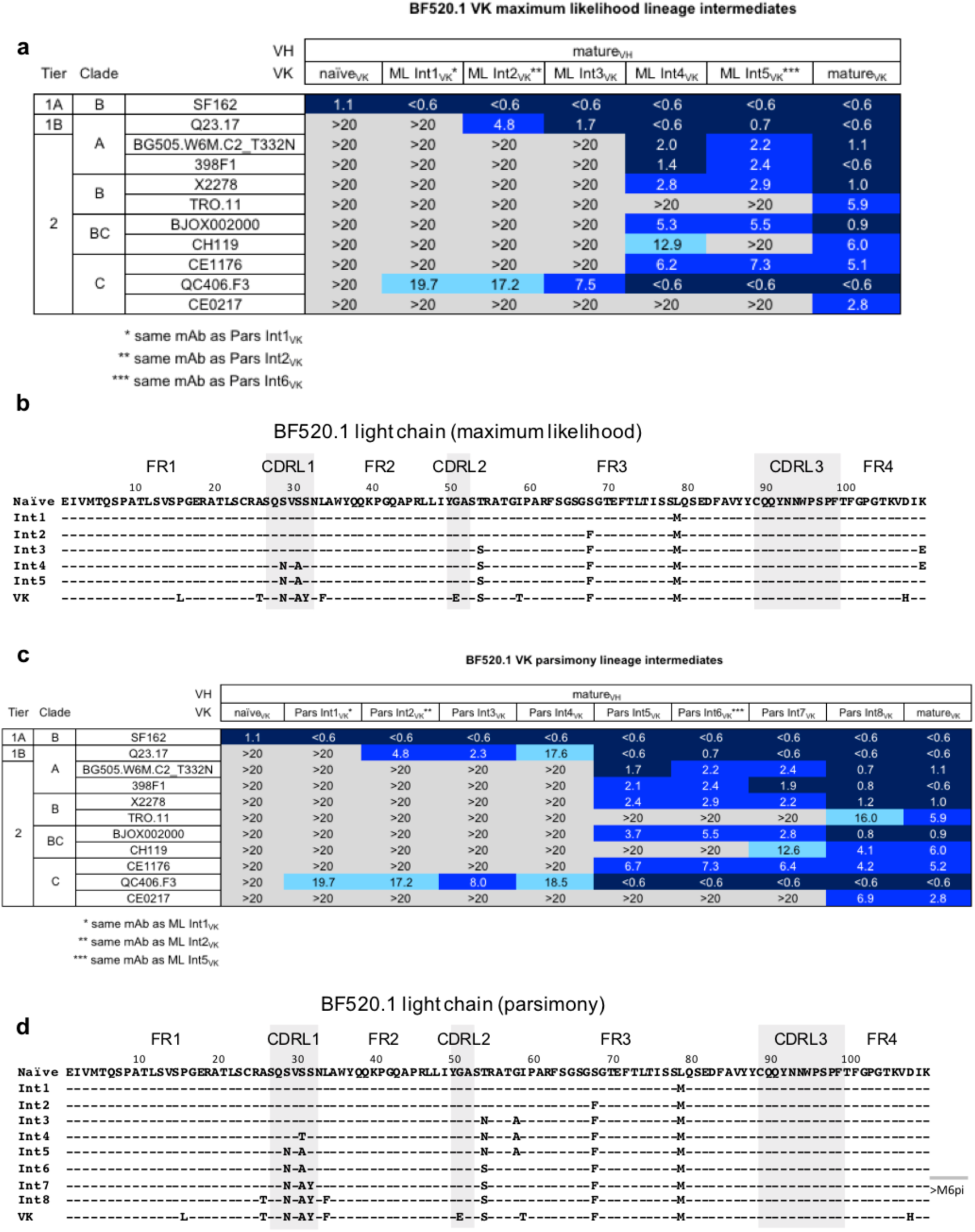
Contribution of kappa light chain maturation to HIV neutralization. BF520.1. a,c, Maximum likelihood (**a**) and parsimony (**c**) lineage kappa chains paired with the mature heavy chain, mAb neutralization of viruses. IC_50_ values (μg ml^-1^) are an average of two to three independent experiments performed in duplicate. IC_50_ values are color-coded with darker shades of blue indicating more potent neutralization. Grey indicates that 50% neutralization was not achieved at the highest mAb concentration tested. **b**,**d**, Amino acid alignments of maximum likelihood (**b**) and parsimony (**d**) kappa chain lineage intermediates. Intermediates designated as >M6 pi were rationally inferred.

**Supplementary Fig. 4:**
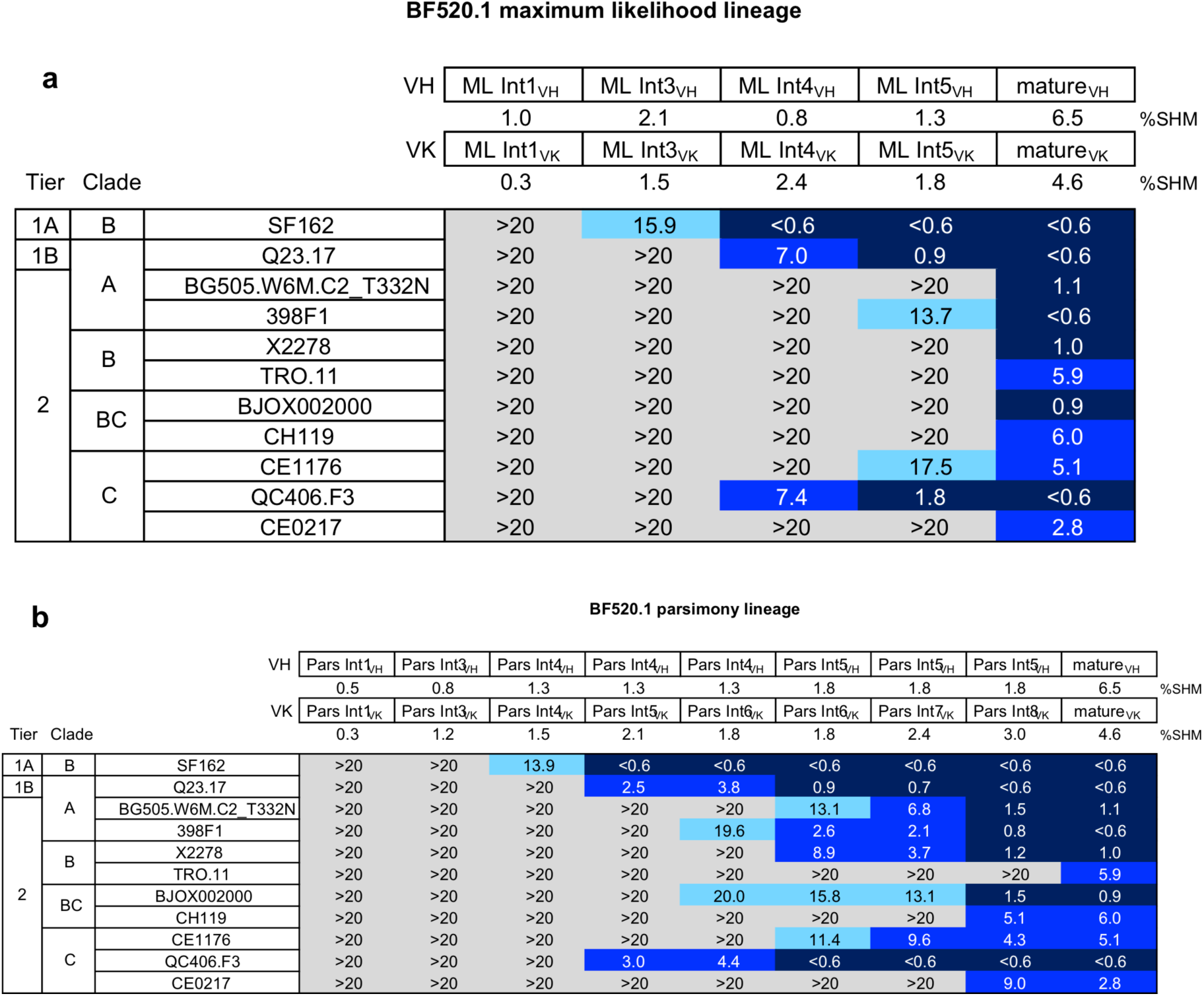
Neutralization properties of paired intermediates of VH and VK. Neutralization of panel viruses by BF520.1 maximum likelihood (**a**) and parsimony (**b**) lineage heavy and light chain paired intermediates. IC_50_ values in μg ml^-1^.

**Supplementary Fig. 5:**
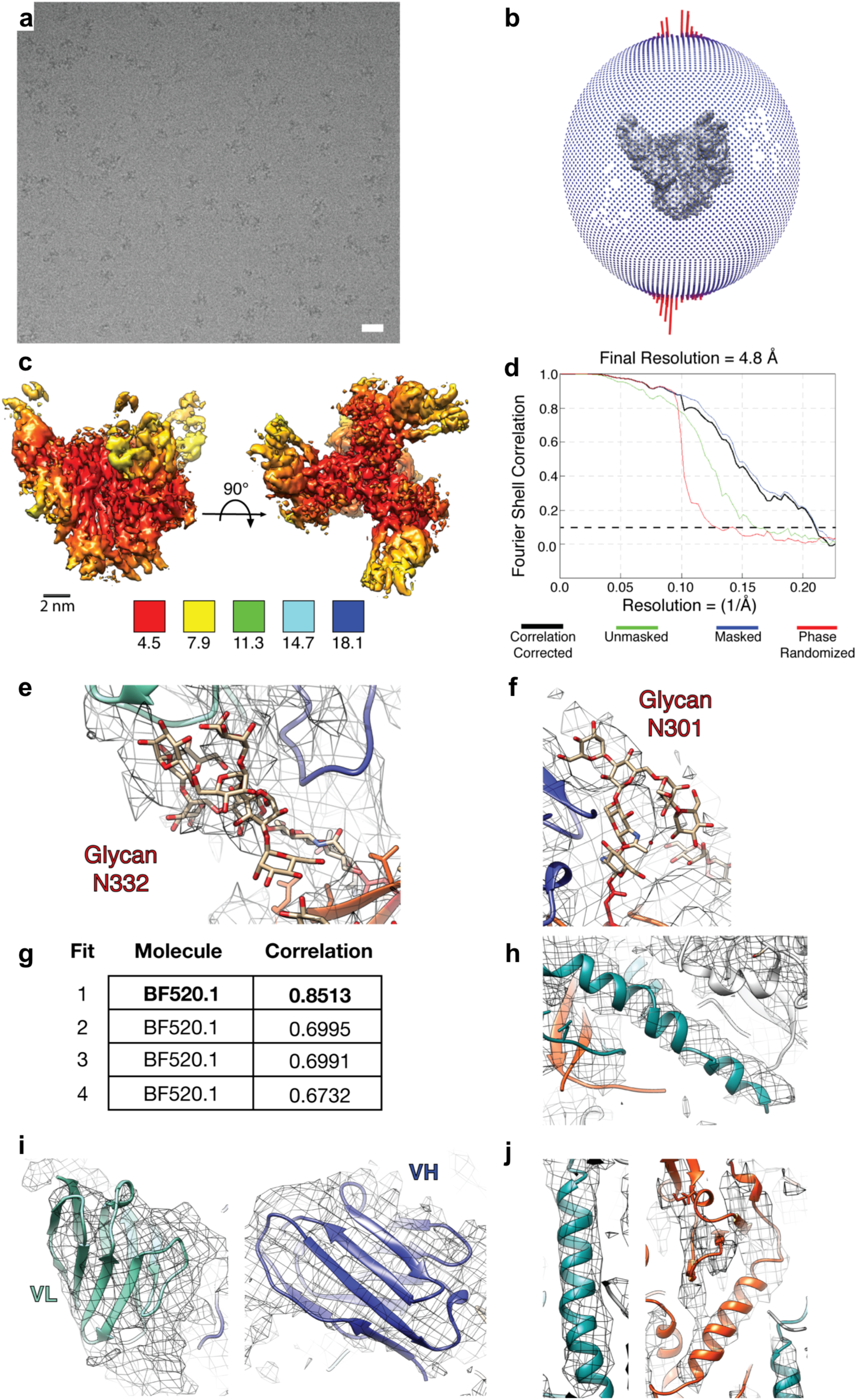
Data collection and refinement of BG505.SOSIP.664 in complex with BF520.1 Fab. a, A sample, frame aligned, micrograph illustrating particle density and distribution. **b**, Angular distribution plot generated in Relion/2.1. **c**, Local resolution analysis of the 4.8 Å cryo-EM reconstruction. The color key shown below indicates local resolution in Å. **d**, Fourier-shell correlation curves for correlation corrected (black), unmasked (green), masked (blue) and phase randomized (red). Dashed line represents the “gold-standard” FSC cutoff of 0.143. **e**,**f**, Densities corresponding to N332 and N301 glycans and positioning of glycans into the cryo-EM density. **g**, Correlation scores of the top fits for the BF520.1 Fab structure into the cryo-EM map. Scores were generated by Chimera UCSF fitmap command utilizing a global search. **h**, Positioning of gp41 HR2 *α*-helix into the cryo-EM density. **i**, Best fit of BF520.1 homology model determined in (**h**). Shown are the variable light chain (left) and variable heavy chain (right). **j**, Positioning of the g41 HR1 *α*-helix (left) and the gp120 *α*1 helix into the cryo-EM density.

**Supplementary Fig. 6:**
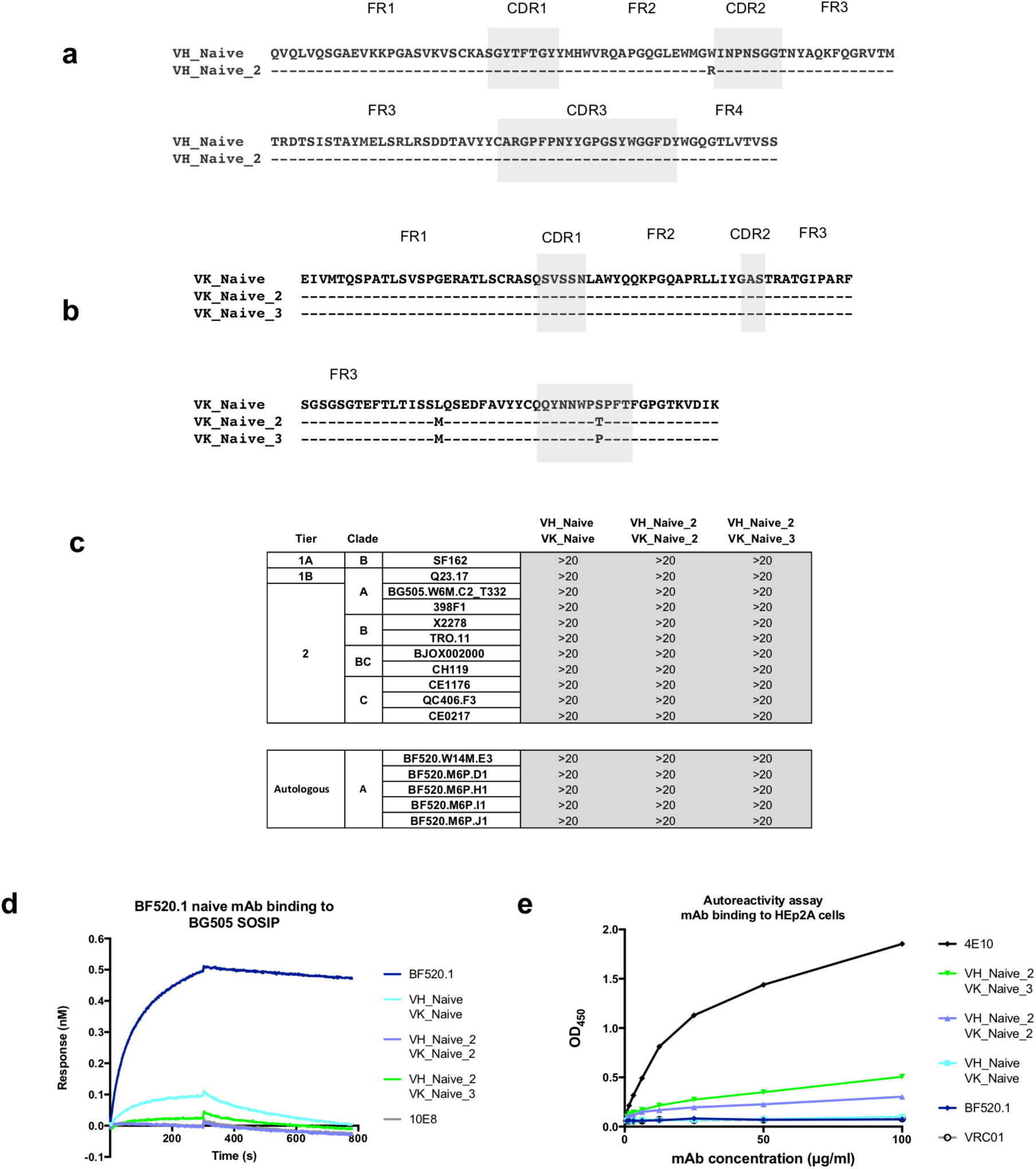
BF520.1 inferred naïve mAbs. a,b, Amino acid alignment of heavy chain (**a**) and kappa chain (**b**) inferred naïve sequences. **c**, BF520.1 naïve mAb neutralization of viruses. IC_50_ values (μg ml^-1^) are an average of two to three independent experiments performed in duplicate. Grey indicates that 50% neutralization was not achieved at the highest mAb concentration tested. **d**, mAb (analyte) binding to the BG505.SOSIP.664 (ligand) measured by BLI. 10E8 is a negative control as the SOSIP trimer does not contain the targeted MPER epitope. Data are representative of two independent experiments. **e**, mAb binding to HEp2A cells measured by ELISA (AESKULISA ANA-HEp-2; Aesku Diagnostics). The MPER-directed bnAb 4E10 is known to be autoreactive and was included as a positive control. VRC01 was included as a negative control. Data are representative of two independent experiments.

**Supplementary Fig. 7:**
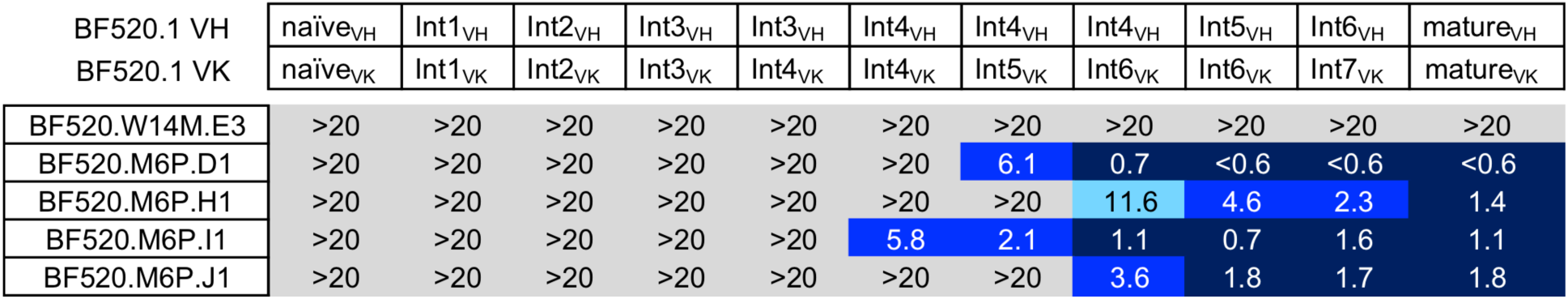
Neutralization of autologous virus by BF520.1 paired lineage intermediates. The origin of the VH and VK chains are shown at the top and are described in Fig. 2b and 3d. Viruses tested are shown to the left and include a variant at first HIV detection (14 weeks of age) designated ‘‘W14’’ and four variants from ∼2.2 months pi (6 months of age, “M6”). IC_50_ values (μg ml^-1^) are color coded with darker shading indicating greater neutralization potency. Results are an average of 2 independent experiments.

## Methods

### Human subject

Peripheral blood mononuclear cell (PBMC) samples were obtained from infant BF520 enrolled in the Nairobi Breastfeeding Clinical Trial^32^, which was conducted prior to the use of antiretrovirals for prevention of mother-to-child transmission. Approval to conduct the Nairobi Breastfeeding Clinical Trial was provided by the ethical review committee of the Kenyatta National Hospital Institutional Review Board, the Fred Hutchinson Cancer Research Center Institutional Review Board, and the University of Washington Institutional Review Board.

### Sample preparation and RNA isolation

PBMCs stored in liquid nitrogen for ∼20 years were thawed at 37°C, diluted 10-fold in pre-warmed RPMI and centrifuged for 10 min at 300x*g*. Cells were washed once in phosphate-buffered saline, counted with trypan blue, centrifuged again, and total RNA was extracted from PBMCs using the AllPrep DNA/RNA Mini Kit (Qiagen), according to the manufacturer’s recommended protocol. RNA was stored at −80°C until library preparation.

We performed library preparation, sequence analysis, and antibody lineage reconstruction in technical duplicate, using the same RNA isolated from each time-point, week 1 (W1) and month 9 (M9).

### Antibody gene deep sequencing

Antibody sequencing was performed as described^46^. Briefly, RACE-ready cDNA synthesis was performed using the SMARTer RACE 5’/3’ Kit (Takara Bio USA) using primers with specificity to IgG, IgK and IgL. The cDNA was diluted in Tricine-EDTA according to the manufacturer’s recommended protocol. First-round Ig-encoding sequence amplification (20 cycles) was performed using Q5 High-Fidelity Master Mix (New England BioLabs) and nested gene-specific primers (Table S1). Amplicons were directly used as templates for MiSeq adaption by second-round PCR amplification (20 cycles). Amplicons were then purified and analyzed by gel electrophoresis and indexed using Nextera XT P5 and P7 index sequences for Illumina sequencing according to the manufacturer’s instructions (10 cycles). Gel-purified, indexed libraries were quantitated using the KAPA library quantification kit (Kapa Biosystems) performed on an Applied Biosystems 7500 Fast real-time PCR machine.

Libraries were denatured and loaded onto Illumina 600-cycle V3 cartridges, according to the manufacturer’s suggested workflow.

**Supplementary Table 1.**
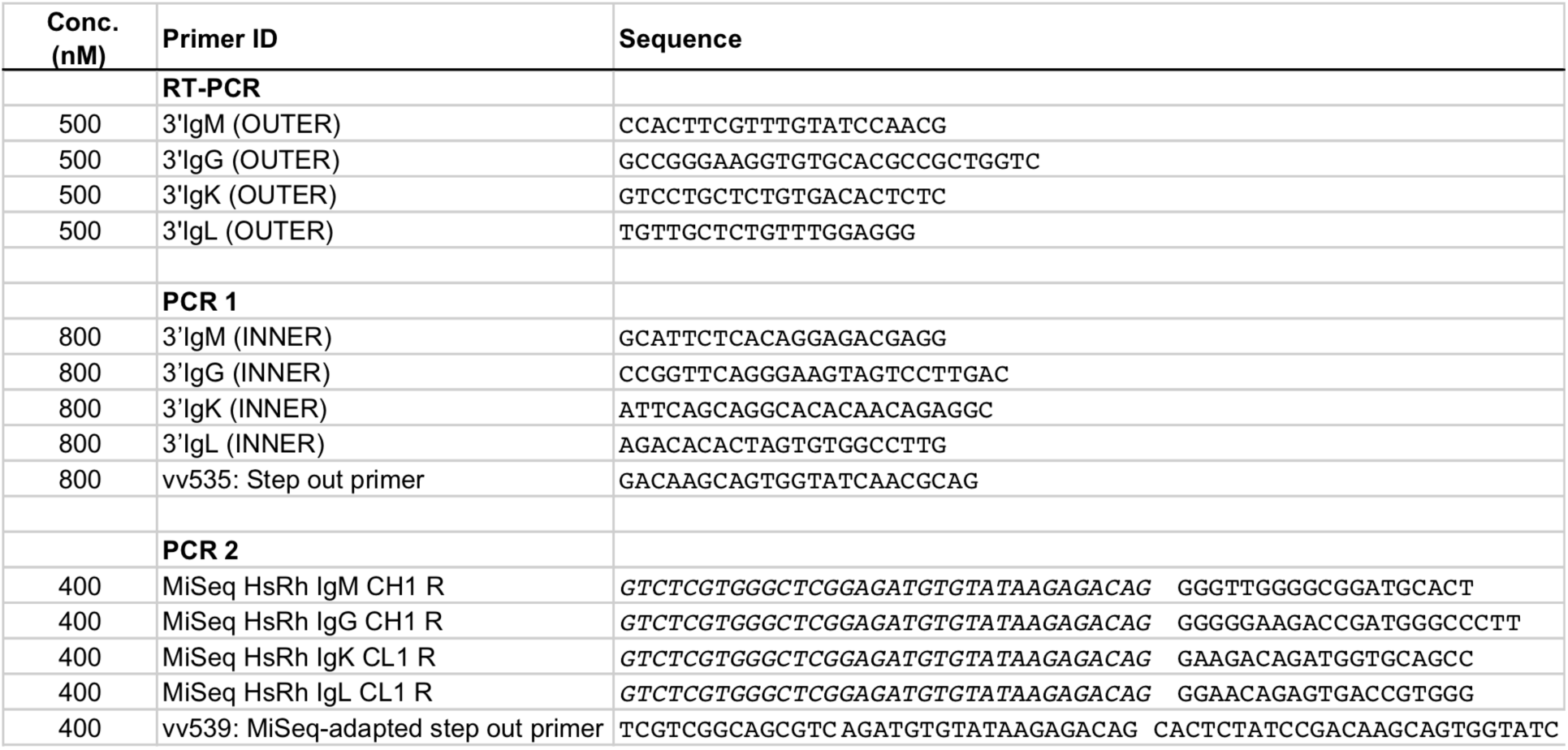
Illumina Miseq library preparation primers.

### Sequence analysis and clonal family clustering

Sequences were preprocessed as previously described^46^. Briefly, amplicons were reconstructed from forward and reverse MiSeq reads using FLASH^47^ and the amplification primers were trimmed using cutadapt^48^. Sequences that contained low-confidence base calls (N’s) were removed (FASTX-toolkit). Filtered sequences from both time-points (W1 and M9) were combined and annotated with partis (https://github.com/psathyrella/partis) using the default options (including per-sample germline inference, Ralph in review at PLOS Comp Biol http://arxiv.org/abs/1711.05843). Sequences with internal stop codons, or with CDR3 regions out of frame were removed during this step. Sequences were then clustered into clonal families using the seed clustering method^22,23^ and the previously-identified BF520.1-IgG and BF520.1-IgK sequences^19^ as “seeds”. This method included the inference of the unmutated common ancestor sequence for each clonal family.

### Antibody lineage reconstruction

Initial phylogenetic trees of BF520.1 heavy and light chain clonal families were inferred using FastTree^49^ and pared down to 100 sequences based on proximity to nodes within the “seed lineage” extending from the inferred naïve ancestor (root) to the mature BF520.1 heavy or light sequence (prune.py, https://git.io/vA7wH). The prune.py program cycles along the path from seed to naive ancestor, adding closest sequences to that path until it achieves the target number of sequences. The 100 clonal sequences were then analyzed by BEAST^26^. BEAST output was summarized for internal node sequences using the approach presented by Gong et al.^25^. In the resulting summary graphic, each red oval represents a unique inferred sequence, with color intensity proportional to the relative confidence that the true lineage included that intermediate. Arrows correspond to amino acid substitutions, with color intensity proportional to the relative confidence that this specific substitution occurred. The most probable BF520.1-IgG lineage paths were consistent between both technical replicates. For BF520.1-IgK, replicates agreed on probable lineage paths but each replicate offered higher resolution at a different part of the lineage (early mutations vs. later mutations). These data were combined to estimate the M0-M9 BF520.1-IgK lineage.

In addition to Bayesian lineage analysis, maximum likelihood (dnaml) and maximum parsimony (dnapars) phylogenetic trees were inferred from the list of 100 clonal sequences, including ancestral sequence reconstruction using PHYLIP tools (http://evolution.genetics.washington.edu/phylip/getme-new1.html) (Felsenstein, J. 2017. PHYLIP (Phylogeny Inference Package) version 3.696. *Distributed by the author. Department of Genome Sciences, University of Washington, Seattle*). Trees were rooted on the inferred naïve sequence and we identified intermediate sequences that were found on lineage paths between naïve to seed sequence. For dnaml and dnapars analyses, filtered IgK sequences were down sampled to 100,000 randomly-selected sequences prior to partis seed partitioning and FastTree trees were pared down using an earlier version of prune.py (https://git.io/vA7ww).

### Validation of Bayesian lineage reconstruction

To computationally validate the performance of Bayesian ancestral sequence inference in antibody lineage reconstruction, we simulated 10 independent clonal families that had similar characteristics to the BF520.1 clonal family and performed Bayesian inference as described in this study on these simulated clonal families. Even though we exclusively used BEAST to perform Bayesian ancestral sequence inference on the BF520.1 clonal family, we additionally performed validation experiments using another popular Bayesian inference software package RevBayes^27^. RevBayes outputs trees that have unconstrained branch lengths, unlike BEAST, which outputs trees with fixed sampling time(s). Clonal family simulation was performed using the procedure described in Davidsen and Matsen (2018)^50^. In short, this procedure approximates germinal center dynamics via nucleotide-context-sensitive mutation and amino-acid selection components and intermediate sampling times. We adjusted the simulation parameters such that the simulation produced clonal families similar to the BF520.1 family in terms of the distribution of root-to-tip distances. To do so, we computed the distribution of the median root-to-tip branch length distances from 10,000 independent RevBayes tree samples for all sequences in the BF520.1 family and matched this distribution with the corresponding distributions from our simulated clonal families (Supplementary Fig. 1b). The simulated “seed” sequence (meant to mimic a sequence of particular interest) was chosen to be the sequence farthest from the root naive sequence.

To assess the accuracy of BEAST and RevBayes in antibody lineage inference, reconstructed ancestral sequences (on the lineage from the naive sequence to the simulated “seed” sequence) were compared to sequences present on the known “true” lineage ancestral sequences in simulated phylogenies. We also ran dnaml and dnapars on these simulated families to compare accuracy between these four methods (Supplementary Fig. 1a). For the Bayesian methods, the posterior probability of a sequence (which we call P(ASR) here for short) is calculated as the number of occurrences of that sequence divided by the number of posterior samples. Any true-lineage sequence that didn’t appear in the posterior was assigned P(ASR)=0. For ML and parsimony, if a sequence was found in the reconstruction, P(ASR) was taken to be 1, and any true-lineage sequence not in the reconstruction got P(ASR)=0. We repeated simulation 10 times with the same parameters, chosen to mimic the BF520.1 clonal family as described above. The results were then collated across simulations to form a single accuracy table, (Supplementary Fig. 1a). Thus, N is the total number of unique reconstructed and “true” lineage sequences across the simulations for a given method.

Our analysis pipeline for the Bayesian phylogenetics component of this work is available at https://github.com/matsengrp/ecgtheow for the purpose of reproducibility. We are currently working on a software tool to enable a more sophisticated version of the analysis presented here.

### VH and VK Lineage Intermediate Pairing

M0-M6 pi intermediates (Bayesian lineage Ints1-3_VH_ and Ints1-4_VK_, Fig. 2b and 3d) were paired together using percentage SHM to assign probable pairings. The remaining post-M6 pi intermediates were paired based on increasing SHM and their incorporation of additional CDR mutations, CDR-adjacent FR mutations and non-conservative FR mutations. Pars and ML lineage intermediates were also paired using percentage SHM to assign probable pairings. Because the M6-M12 pi rationally inferred substitutions would be the same for ML and Pars lineages, these additional changes were added to the Pars lineage only.

### mAb Preparation

Antibody heavy and light chain variable regions were synthesized as “gBlocks” by Integrated DNA Technologies (www.idtdna.com) and subsequently cloned into IgG and IgK expression vectors as previously described^51^. Equal ratios of heavy and light chain plasmids were co-transfected into 293F cells using FreeStyle MAX (Invitrogen) according to the manufacturer’s instructions. Protein G columns were used to purify IgG as previously described^52^.

### Pseudovirus Production and Neutralization Assays

Pseudovirus production and neutralization assays were performed as previously described^53^ with an alternative cell lysis and B-galactosidase detection system (Gal-Screen; ThermoFisher). Specifically, 85uL of the 150μL total volume was removed from each well (50μL remaining) and 50μL of Gal-Screen substrate (diluted 1:25 in “Buffer A”) was added to each well. Luminescence was measured after 40 minutes incubation at room temperature. The mAbs were diluted 2-fold from 20 to 0.6 μg ml^-1^. IC_50_ values represent the concentration (μg ml^-1^) at which 50% of the virus was neutralized and are the average of two or three independent experiments performed in duplicate.

### Biolayer Interferometry Assays

Binding of mAbs to HIV Env SOSIP trimers was measured using biolayer interferometry on an Octet RED instrument (ForteBio). For the qualitative comparison of lineage intermediate binding, IgG antibodies diluted to 10 μg ml^-1^in PBS plus 1% BSA, 0.01% TWEEN-20, and 0.02% sodium azide were immobilized onto anti-human IgG Fc capture (AHC) biosensors and BG505.SOSIP.664 (diluted to 1μM in the same buffer) was flowed as analyte in solution. For mAb kinetic determination, BG505.SOSIP.664-AviB (25 μg ml^-1^) was immobilized onto Streptavidin (SA) biosensors. Varying concentrations of IgG were flowed as analyte in solution. A series of six, two-fold dilutions of naïve_VH_mature_VK_ and mature_VH_naïve_VK_ mAbs (667 to 21 nM) as well as 0 nM IgG were tested. The BF520.1 naïve mAb concentrations ranged from 2.67 to 0.33 μM (series of four, two-fold dilutions) and 0 μM. Association was monitored for 6 minutes and dissociation for 10 minutes. Binding-affinity constants (K_D_; on-rate, K_on_; off-rate, k_dis_) were determined using ForteBio’s Data Analysis 7.0. Average measurements from reference wells were subtracted and data were processed by Savitzky-Golay filtering prior to fitting using a 1:2 (bivalent analyte) model of binding.

### Sample preparation for cryo electron microscopy

Purified BG505.SOSIP.664 trimers^54^ were mixed with 4-fold molar excess BF520.1 Fab and diluted to 0.4 mg/mL in PBS or PBS supplemented with 70 µM n-Dodecyl-*β*-D-Maltoside (DDM). The mixture incubated for 1 hour at room temperature prior to vitrification. A 3.0 µL aliquot was applied at 4°C and 100% humidity to glow-discharged C-Flat 1.2/1.3 4C holey carbon-coated grids (Electron Microscopy Sciences), blotted, and immediately plunge frozen in liquid ethane by using a Vitrobot Mark IV specimen preparation unit (FEI Co.).

### Cryo-EM data collection

Vitrified grids were imaged using an FEI Titan Krios operating at 300 keV and equipped with a Gatan K2 summit direct detector device. Micrographs were collected at 130,000×, corresponding to a pixel size of 0.55 Å/pixel in super resolution mode. Each image received a dose rate of ∼ 8 e^-^/pix/s with 200 ms exposure per frame, and an estimated defocus ranging from 1.0 - 3.5 µm. Data were collected in three separate sessions, resulting in a total of 3454 images using EPU (908 micrographs) (FEI) and Leginon (2546 micrographs)^55^ automated data collection softwares.

### Cryo-EM data processing

Frame alignment and CTF estimation were carried out independently for data collected using EPU and data collected using Leginon softwares. Frame alignment and dose-weighting were completed using MotionCor2^56^, and CTF estimation was performed using CTFFIND4^57^. Relion 2.1^58^ was utilized for further processing and 3D refinement. Approximately 1,000 particles were manually selected and subjected to 2D classification to build templates for automated particle picking. A total of 559,022 particles was selected and binned to 8.8Å/pixel for expeditious processing. The binned particle stack was subjected to 2D classification where 116,972 were selected for 3D refinement. This particle stack was re-extracted as a 4× binned stack (pixel size 2.2Å/pix). Approximately 32,000 particles were chosen for 3D classification and a subset of ∼9,000 particles was selected to build a low resolution 3D model, resulting in a 9.02Å map. This model was low-pass filtered to 60Å and refinement was performed on the full stack of 116,972 particles in Relion 2.1 with C3 symmetry imposed. Map sharpening and post processing in Relion yielded a 4.8Å structure using the “gold-standard” FSC cutoff of 0.143.

### Model Building

The atomic model was generated by first fitting the BG505.SOSIP.664 trimer from the BG505:PGT128 cryo-EM structure (PDB ID: 5ACO)^34^, with glycans temporarily removed, into the generated 4.8Å map using the fitmap command in UCSF Chimera^59^. Overall, docking of the BG505 structure showed a good quality fit resulting in a correlation score of 0.8602. No further modification to the protein structure was performed. Both glycans at positions N332 and N301 were manually placed into their corresponding densities in the 4.8Å map.

The BF520.1 variable heavy and light chains were submitted as a single polypeptide sequence to MUlti-Sources ThreadER (MUSTER) Online to predict the structure of BF520.1^60^. The structure template used for structure determination was a single chain variant of anti-gp120 antibody, b12 (PDB ID:3JUY)^61^. Using Chimera’s fitmap command, the BG505.SOSIP.664 trimer and BF520.1 variable domain were sequentially docked into the 4.8Å map.

